# Development of transformation for genome editing of an emerging model organism

**DOI:** 10.1101/2022.04.19.488772

**Authors:** Yutaka Yamamoto, Susan A. Gerbi

**Affiliations:** Department of Molecular Biology, Cell Biology, and Biochemistry Brown University Division of Biology and Medicine Sidney Frank Hall, 185 Meeting Street Providence, RI 02912, USA

**Keywords:** piggyBac vector, fungus fly *Sciara (Bradysia) coprophila*, germline transformation

## Abstract

With the advances in genomic sequencing, many organisms with novel biological properties are ripe for use as emerging model organisms. However, to make full use of them, transformation methods need to be developed to permit genome editing. Here, we present development of transformation for the fungus fly *Sciara (Bradysia) coprophila*; this may serve as a paradigm for development of transformation for other emerging systems, especially insects. *Sciara* has a variety of unique biological features including locus-specific developmentally-regulated DNA amplification; chromosome imprinting; a monopolar spindle in male meiosis I; non-disjunction of the X chromosome in male meiosis II; X chromosome elimination in early embryogenesis; germ line limited (L) chromosomes; high resistance to radiation. Mining the unique biology of *Sciara* requires a transformation system to test mutations of DNA sequences that may play roles for these features. We describe a *Sciara* transformation system using a modified piggyBac transformation vector and detailed protocols we have developed to accommodate *Sciara*-specific requirements. This advance will provide a platform for us and others in the growing *Sciara* community to take advantage of this unique biological system. In addition, the versatile piggyBac vectors described here and transformation methods will be useful for other emerging model systems.

**Author Biographies:** <u>Susan A. Gerbi</u> (Ph.D. with Joseph Gall at Yale University 1970) is the George Eggleston, Professor of Biochemistry at Brown University. Her research includes chromosomes, DNA replication and ribosomal RNA. She was President and is a Fellow of ASCB, a Fellow of AAAS and received the RI Governor’s Award for Scientific Achievement. Other honors include RNA Society/CSHL Press Distinguished Research Mentor award; GSA George Beadle award; ASCB Senior Leadership/ Mentoring Award. She is a national leader in graduate education, including member of the National Academy of Sciences Panel on Bridges to Independence that led to the NIH K99 program, Chair of the AAMC Graduate Research Education Training Group; Chair of the FASEB Consensus Conference on Graduate Education. <u>Yutaka Yamamoto</u> (M.D. Kansai Medical University 1990; Ph.D. with Walter Gehring at Biozentrum – Basel 1995; postdoc with David Glover at Dundee University and University of Cambridge) is a research associate at Brown University.

**Graphical Abstract:** 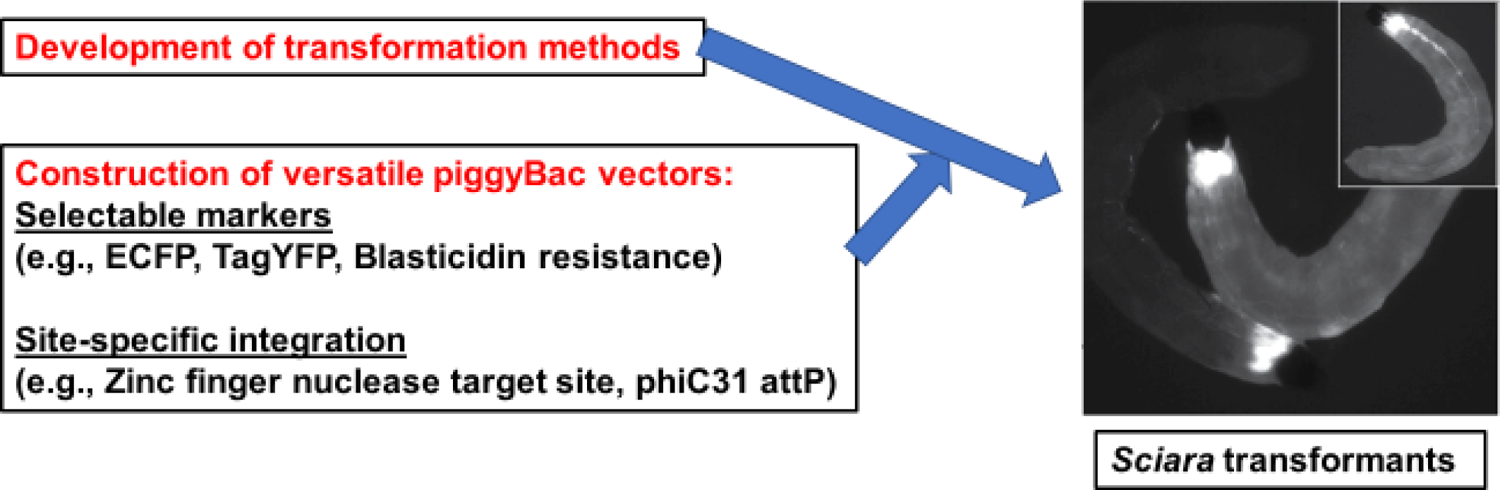

## 1. Introduction

Many advances in biological understanding have come from elucidation of principles in unique biological systems. For example, the protozoan ciliate *Tetrahymena* served as the experimental material for discoveries that lead to two different Nobel prizes (self-splicing, telomerase)(Pederson 2010). Similarly the nematode worm *C. elegans* has been the experimental organism that has garnered several Nobel prizes (programmed cell death, RNA interference). Recognition of the value of the breadth of knowledge from a large number of model systems (Sullivan 2015) has led to efforts to sequence genomes from a diverse array of organisms, including the i5K project to sequence the genomes of 5000 arthropods including all phylogenetic branches of insects (http://arthropodgenomes.org/wiki/i5K). In order to reap the full advantage of such information, transformation methodology is needed that has broad applicability. The recent advent of CRISPR/Cas9 has provided a technology for genomic mutations (Jinek et al. 2012; Mali et al. 2013; Esvelt et al. 2014). However, since Cas9 cleavage creates blunt ends, it is not suitable for insertion of large DNA (such as genes) using the non-homologous end-joining pathway that is preferred in most organisms over homologous recombination. In contrast, piggyBac with its broad phylogenetic range allows a widely adaptable means for genomic insertion of large DNA. Here we present the first report of a method using piggyBac to transform the fungus fly *Sciara coprophila*, with no prior transformation system, and this may offer applications to other emerging model organisms, especially insects.

The fungus fly *Sciara* (Meigen 1803), represents the largest known eukaryotic genus in the world, with over 700 species. Some years ago, it was renamed *Bradysia coprophila* (Steffan, 1966), but we retain the earlier name of *Sciara* to be consistent with the large body of published literature on this organism. The novel biological strategies employed by *Sciara* led it to be on the short list as a premier model organism in the 1930s at a Cold Spring Harbor meeting when *Sciara* competed with *Drosophila melanogaster* and in the 1970s when Sydney Brenner compared the merits of *Sciara* with the worm *Caenorhabditis elegans*. The many unique features of *Sciara* (reviewed by Gerbi, 1986; Gerbi and Urnov, 1996; Gerbi et al., 2002; Gerbi 2022) include:

- Locus-specific DNA amplification in larval salivary gland polytene chromosomes;
- chromosome imprinting;
- a monopolar spindle in male meiosis I;
- non-disjunction of the X chromosome in male meiosis II;
- chromosome elimination in early embryogenesis;
- germ line limited (L) chromosomes;
- high resistance to radiation.

Recently, the *Sciara* genome has been sequenced, annotated and assembled (Urban et al 2021). This serves as the foundation for genome editing where a transformation system for *Sciara* is required to facilitate experiments to understand the molecular basis for this extraordinary number of unique biological features. Such molecular tools would allow manipulation of the *Sciara* genome to assay the role of specific DNA sequences in these unique biological features.

Historically, transposable elements have permitted integration of DNA sequences into insect genomes. P-elements were the first of these (Rubin and Spradling 1982), but they lacked a broad host range (Handler et al. 1993; O’Brochta and Atkinson 1996; Handler et al. 1998). In contrast, piggyBac has been used to transform many organisms, including a broad range of insects from Diptera, Lepidoptera and Coeoptera (reviewed by Handler 2002; Sarkar et al. 2003). Moreover, P-elements preferentially insert into the promoters of genes, whereas piggyBac has evolved to reduce deleterious effects in host gene expression (Bellen et al. 2011). The piggyBac transposon, derived from the cabbage looper moth *Trichoplusia ni*, is a 2472 bp Class II transposon with 13 bp inverted terminal repeats (Cary et al. 1989; Fraser et al. 1996). PiggyBac contains a single ORF that encodes a 594 amino acid transposase that mediates insertion of this transposon into a TTAA target site in the genome, resulting in target-site duplication that flanks the transposable element (Fraser et al. 1995 and 1996). The piggyBac transposase is active in a broad number of insects (Handler 2002) and other animals, ranging from planaria to mammalian cells (Lobo et al. 2006).

Given the extensive host range of piggyBac, we have developed a transformation system for *Sciara* based on the piggyBac transformation vector pBac[3xP3-ECFPafm] (Horn and Wimmer 2000; Horn et al. 2000), which is designed to screen transgenic animals using the ECFP fluorescent marker driven by the highly conserved P3 promoter. Our subsequent modifications included use of TagYFP and blasticidin resistance as markers for screening. For future versatility to allow site specific insertions of large DNA by a variety of means, we included an artificial Zinc Finger Nuclease Target (ZFN-T) site (Urnov et al. 2005) and the phiC31 attP site (Groth et al. 2004). We also report here the development of methods for the various steps in *Sciara* transformation. These vectors and procedures will serve as the foundation for future experiments on *Sciara* by the growing scientific community. Also, they should be helpful for transformation of other non-model organisms.

## 2. Materials and Methods

### 2.1. Sciara injection

#### 2.1.1. Egg collection and injection

The sterilized vials used for rearing the fungus gnat *Sciara coprophila* (syn. *Bradysia coprophila* Lintner) are 29 mm. inner diameter and 9.5 cm tall (Fisher # 03-339-26H or Wilmad LabGlass), filled with 1.3 inches of 2.2% (wt/vol) bacteriological agar and topped with non-absorbant cotton-filled gauze plugs. One day before injection, 5 vials were set up with mass matings, with each vial containing 10 males and 10 females that are male producers from the HoLo2 line that was derived from the 7298 stock (X’ chromosome for female producers marked with the dominant allele for *Wavy* wings) but lacks the polymorphism noted for 7298 (Yamamoto et al 2021). Previously, the larvae should be well-fed to give fat adults with large eggs from the females. All the following steps for egg collection and transformation were performed in a room at 18^0^C with several humidifiers for at least 70% humidity and preferably >90% humidity. We used three humidifiers in a small temperature controlled room with the recirculating air duct blocked, and we monitored the temperature and humidity level. One day (18 hours) after mating, the adult females were collected by anesthesia on a CO_2_ pad and their thorax is gently squeezed (avoiding injury to the abdomen) to induce their wings to extend upright. They were then picked up by their wings with a forceps for placement on a 100 mm diameter petri dish containing 2.2% (wt/vol) agar, with their wings inserted into the agar. Egg laying was induced by gently squeezing the head with a forceps until the female fly showed seizure-like movements. The petri dish was then covered with the lid that was moistened with a water dampened kimwipe to prevent electrostatic attraction of the eggs to the lid. After 30 minutes the eggs were transferred with a dampened (but not overly wet) fine bristle paint-brush onto the egg sorting agar, gently using forceps to dislodge any embryos stuck on the brush when transferring. It is important to proceed quickly so that injections are done within a window of 1-3 hours after egg-laying, before the pole cells form and while the embryos are still a syncytial blastoderm (DuBois 1932a and 1932b and 1933; de Saint Phalle and Sullivan 1996).

Egg sorting petri plates (50 mm diameter and 9 mm height; Falcon) were made with 2.2% (wt/vol) agar darkened by adding green or blue food dye, so that the nuclei inside the eggs could be visualized with a strong focused light. The colored agar was removed with a metal spatula from the petri dish and placed upside down on top of the lid. A straight edge (about 4 cm long) was made by cutting the agar with a razor blade 1/5 from the edge of the petri dish. Re-usable glass spatulas were made in advance by heating with rotation the long stem of a Pasteur pipette in a Bunsen burner flame, removing it from the flame to pull and break the melted tip, and heat sealing the tip (close, typically 3 cm, to the thicker body of the pipette) in the flame so that the sealed tip is the width of one embryo. The sealed tip is round and not flat. The eggs were aligned close to each other with the glass spatula, with the anterior end closest to the cut edge of the agar. The eggs should not be right at the edge nor protrude beyond the agar cut edge as they will dry out. The posterior part of the embryo is opaque; the anterior part is more transparent and the nuclei can be visualized. Embryos with a small clearing at the anterior and posterior ends should not be used as they are too old.

*Sciara* appeared to be extremely sensitive to the glues commonly used for *Drosophila* embryo injection. Instead, Kwik-Cast glue (World Precision Instruments, Inc) was used as a non-toxic substitute. A small amount of Kwik-Cast yellow and blue substances were squeezed from the tubes and mixed on a glass microscope slide with a pipetteman tip to turn the mixture to green. A dab of the green Kwik-cast mixture was scooped up with a pipetteman tip and spread evenly on the long edge of a fresh, ethanol-washed glass microscope slide. The thickness of the Kwik-Cast is crucial for successful injection. Embryos must be lightly covered with Kwik-Cast.

However, if it is too thick, it will interfere with the injection. Kwik-Cast cures in 3-5 minutes at 18^0^C. Moreover, for successful transfer of the embryos, the viscosity of the Kwik-Cast is important. If it is too soft, the embryos will remain on the agar plate. Conversely, if it is too hard, it will not cover the embryos enough. The viscosity of the Kwik-Cast glue was monitored by touching the Kwik-Cast mixture that remained on the first slide with a pipetteman tip. When the viscosity was ok (∼3 minutes after mixing the Kwik-Cast), the microscope slide with the long line of Kwik-Cast was inverted and held against the embryos that were aligned on the colored agar to transfer them to the Kwik-Cast, with the posterior end of the aligned embryos at the edge of the slide. Any excess glue between the embryos and the edge of the slide can be removed with a razor blade. The embryos on the Kwik-cast glue stripe were then covered lightly with halocarbon 27 oil (CAS#9002-83-9; Halocarbon Products Corporation) and placed in a humid chamber for at least 3 minutes to let the glue cure completely. The humid chamber was a shallow plastic box (245 mm X 245 mm; Corning) with 2-3 layers of 3 MM paper soaked with water; the microscope slides were elevated on plastic spacer bars to avoid direct contact with water. Note that halocarbon 27 oil is a medium weight polymer of chlorotrifluoroethylene (PCTFE), rather than the high molecular weight PCTFE polymer used for *Drosophila* injections.

*Sciara* embryos were injected within 2 hours of egg laying, using an injection apparatus similar to that used for *Drosophila,* with needles drawn out from borosilicate glass capillaries (1B100-4; World Precision Instruments, Inc.) and filled with a Microfil 28 needle (MF28G67-5 4; World Precision Instruments, Inc.). The chorion of *Sciara* is very thin and does not need to be removed for injection; however, the *Sciara* embryos need to be maintained in a humid environment to avoid desiccation. PiggyBac vector DNA (final concentration in injection solution of 1 μg/μl water) was mixed with hr5-ie1 piggyBac transposase plasmid (400 ng/μl) and IE1 transactivator plasmid (pIE1/153) (0.4 ng/μl) for the injection (Mohammed and Coates 2004).

The inverted bright field microscope used for the injections rested on an anti-vibration platform (World Precision Instruments, Inc). The filled needle was placed in the injector and lowered to contact the edge of a broken coverslip that was affixed with double-sided tape to the microscope slide to break the needle tip, allowing droplets of the injection fluid to flow continuously from the tip. The needle was lowered to be in line with the embryos on the microscope slide, and the slide was then moved towards the needle for sequential injection into the posterior end of each embryo. During the injection procedure, the pressure of liquid in the needle was adjusted to maintain a continuous flow; too little pressure results in cytoplasm from the embryo backing up into the needle. Alternatively, pulse injections were used with a time gated setting for the injector.

After injection, the slide with affixed embryos was held vertically for 5 minutes on a paper towel to drain off the excess oil. The slide was examined under a dissecting microscope to confirm that just enough oil was left to cover the eggs by surface tension. If the embryos are completely submerged in too much oil, they will not survive due to lack of oxygen. *Sciara* seems to be very sensitive to the oxygen level. Excess oil was blotted off with a rolled up kimwipe, the slide was placed in a humid chamber box (described above) that was wrapped in Saran wrap and incubated 5 days at 18^0^C.

After 5 days at 18^0^C and development into late embryonic stages (before pigmentation of the jaw appears), the embryos were excised from the Kwik-cast using the tip of a 23G needle to cut the edge of the glue surrounding the embryos. This allowed the oil to seep under the embryos and pop them off the Kwik-cast glue. The embryos were then transferred with a glass spatula onto a 100 mm diameter petri dish containing 2.2% agar (wt/vol) (without antibiotics), and placed in a humid chamber at 21^0^C. It is important to do this before the embryos emerge as first instar larvae are more prone to damage by the transfer procedure than the embryos. Once the larvae emerged, they were fed on alternating days 3X per week. *Sciara* food is one part Brewer’s yeast mixed with 4 parts (by volume) of an autoclaved mixture of 2 parts very finely ground oat straw, 1 part mushroom powder and 1 part spinach or nettle powder (all are insecticide-free). If the larvae try to crawl out of the petri dish (regularly monitor this), the petri dish containing them should be moved from the humid chamber to a dry chamber at 21^0^C. After the 4^th^ instar and pupation, when the body color of the pupae had begun to darken, the pupae were transferred from the petri dish into a glass vial containing 2.2% agar (wt/vol) and incubated at 21^0^C until the adult flies emerged. About 300 *Sciara* embryos were injected in one day to become ∼60 G0 flies, based on the observation that typically there are 20-50% survivors at 5 days after injection, and after emerging as 1^st^ instar larvae at least 60-70% develop into fertile adults. The injection is repeated on sequential days with the goal of obtaining 100 fertile G0 adults that will yield 1-2 transformants.

#### 2.1.2. Subsequent crosses with fluorescent marker selection

When the G0 male flies emerged, each was placed in a vial containing 2.2% agar (wt/vol) with 5 female producer female flies (HoLo 2 line). Alternatively, each male fly was crossed with 3 females on day one and then on day two the male was moved to a fresh vial containing 2 new female producer female flies. Therefore, each single G0 male was crossed with a total of 5 female producer females. The vials were incubated at 21^0^C until the larva were 3^rd^ instar, when the larvae from a single G0 male were transferred to a 100 mm diameter petri dish containing 2.2% agar (wt/vol), with a maximum density of 50 larvae per petri dish. Since food in the larval gut can create autofluorescence, the larvae were left overnight at 18^0^C on the petri dish with no food to clear the gut contents. Larvae with fluorescent signals were selected under a fluorescence stereo dissecting microscope (Zeiss Stemi SV11 Apo, equipped with a standard EYFP, EGFP, ECFP fluorescent filter set and with an X-Cite 120 light source) and fed 3X per week on the agar petri dishes at 21^0^C until they pupated. Fluorescent screening was done as soon as feasible (generally for 3^rd^ rather than 4^th^ instar larvae). After the fluorescent larvae from a single G0 male emerged as pupae and started to show strong eye pigmentation (and the fluorescent signal may no longer be visible), they were transferred to a single vial containing 2.2% agar (wt/vol) for development into adult female flies. To amplify the transformant lines, matings were done of transformed female-producer females with males from the *Sciara* HoLo 2 line, using three males per female in a single vial. Thus, the transgene is transmitted through the female in each generation, avoiding the problem that male-derived chromosomes are eliminated on the monopolar spindle in meiosis I of spermatogenesis (where the transgene could be lost).

#### 2.1.3. Subsequent crosses with antibiotic selection

When the G0 male flies emerged, each was crossed with 5 female-producing female flies (HoLo 2 line) as in section 2.1.2. One day after each cross, the females were placed onto 100 mm diameter petri dishes containing 2.2% agar (wt/vol) with 20 µg/ml blasticidin S (A.G. Scientific, Inc), diluted from a stock solution of 10 mg/ml blasticidin in 20 mM HEPES, pH 7.2. The petri dishes were incubated in a humid chamber at 21^0^C, with removal of the dead adult females after 2-4 days. As a control, non-transformed female-producing females (HoLo 2 line) were mated with HoLo 2 males for egg laying on petri dishes containing 2.2% agar (wt/vol) with 20 μg/ml blasticidin. The petri dishes with the control and experimental larvae were maintained without feeding in a humid chamber at 21^0^C until the control larvae died (generally 3^rd^ instar). At that point, the blasticidin-resistant larvae were transferred to petri dishes containing 2.2% agar (wt/vol) but no blasticidin and were maintained in a dry chamber at 21^0^C with feeding 3X per week. When the larvae became dark pupae, they were transferred from the petri dishes to vials containing 2.2% agar (wt/vol). When the adult flies emerged, the female producer female flies were crossed with HoLo 2 males to maintain the transgene through the female lineage and to amplify the transformant lines.

### 2.2. Constuction of piggyBac vectors

#### 2.2.1. Construction of pBac[3XP3-ECFPafm] attP-ZFN-T

A 231 bp DNA fragment containing the attP sequence (AATAATGATTTTATTTTGACTGATAGTGACCTGTTCGTTGCAACAAATTGATGAGCAATGCTTTTTTATAATGCCAACTTTGTACAAAAAAGCTGAACGAGAAACGTAAAATGATATAAATATCAATATATTAAATTAGATTTTGCATAAAAAACAGACTACATAATACTGTAAAACACAACATATCCAGTCACTATGAATCAACTACTTAGATGGTATTAGTGACCTGTA) with PstI sites flanking each end was synthesized by Integrated DNA Technologies (IDT) and cloned into the PstI site of pBac[3XP3-ECFPafm] attP (Horn and Wimmer 2000) that was a kind gift from Dr. Alfred M. Handler, USDA. The synthesized fragment was cloned into the pCR-Blunt II-TOPO vector (Invitrogen).

Next, a linker was added that included the same artificial zinc finger nuclease (ZFN) target site (CGCTACCCGACCATGAAGCAGCA) as used by Urnov et al. (2005). The FseI sites flanking the linker were created by annealing two oligos (FTEAF 5’and FTEAF 3’). The sequences of these oligos were

FTEAF 5’: CCCGCTACCCCGACCATGAAGCAGCAGAATTCGGCGCGCCGGCCGG;

FTEAF 3’: CCGGCGCGCCGAATTCTGCTGCTTCATGGTCGGGGTAGCGGGCCGG.

This FseI bounded linker containing the ZFN target site was then cloned into the FseI site of (pBac[3XP3-ECFPafm] attP) to create (pBac[3XP3-ECFPafm] attP-ZFN-T).

#### 2.2.2. Construction of pBac[3XP3-TagYFP, hr5-ie1-BlasR, su(Hw)BS] attP-ZFN-T

Since the introduction of TagYFP involved PstI restriction digestion, attP was initially removed from the PstI site and later re-introduced into that site. The removal of attP converted the plasmid pBac[3XP3-ECFPafm] attP-ZFN-T described in the preceding section (2.2.1) to pBac[3XP3-ECFPafm] ZFN-T as the starting vector. As described in detail below (also see Supplementary Figure S3), two su(Hw)BS were introduced, then ECFP was replaced with TagYFP, followed by addition of the attP site.

The suppressor of Hairy-wing binding site (su(Hw)BS) was introduced into the plasmid in a two-step procedure as follows. First, the plasmid pBac[3xP3-ECFP afm] ZFN-T was linearized by PCR using the primer sets pBac 3Fp and pBac 3R:

pBac 3Fp: phos-GATGTTCCCACTGGCCTGGAGCG

pBac 3R: GATGTTTTGTTTTGACGGACCCCTT

The 362 bp sequence for su(Hw)BS was amplified from the pYES vector (Patton et al. 1992)(kindly provided by Dr. John Tower, University of Southern California) using primer sets 20-1-1 F5p and YY59-1:

20-1-1-F5p: phos-AGTTACTCTTCCGGCTAAATGGTATGGCAAGAAAAGGTATGCAATAT

YY59-1: GGGCGAATTGGGTACCCTATTCGC

The amplified su(Hw)BS sequence was then blunt-end ligated into the linearized vector pBac[3xP3-ECFP afm] ZFN-T noted above at a position just in front of 3XP3-ECFP. This clone was subsequently linearized by PCR using primer pairs pBac 5F and pBac5Rp:

pBac 5F: GACAATGTTCAGTGCAGAGACTCG

pBac 5Rp: phos-AGATCTGTCATGATGATCATTGCAATT

Subsequently, the same amplified su(Hw)BS sequence noted above was again blunt ligated, in this case just downstream of ZFN-T in the plasmid being constructed.

Next, the ECFP gene was replaced by TagYFP in the plasmid under construction, named pBac[3XP3-ECFP, su(Hw)BS] ZFN-T. The 750 bp TagYFP coding sequence was amplified from pTagYFP-C (Evrogen) using primer pairs P3-TagYFP F1 and TagYFP 3R1:

P3-TagYFP F1: gcccgggatccaccggtcgccaccATGGTTAGCAAAGGCGAGGAGCTGTTCGCCGGC

(The TagYFP coding sequence is underlined and the 3xP3 is in lower case letters)

TagYFP 3R1:

AGAGTC<u>GCGGCCGC</u>TTTACCGGTACAGCTCGTCCATGCCGTGGGTGTGGC

(The NotI site is underlined)

Subsequently, the 1.4 kb fragment including (including the 5’ piggyBac arm and 3XP3-TATA) was amplified from the plasmid noted above, named pBac[3XP3-ECFP, su(Hw)BS] ZFN-T, using primer pairs PstI F1 and P3-TagYFP R1:

PstI F1: CCTA<u>CTGCAG</u>GTCATCACAGAACACATTTGGTCTAGCGTGTCCACTCCGCC

(The PstI site is underlined)

P3-TagYFP R1: (contains 3’ end of P3 promoter and 5’ end of TagYFP coding sequence) GGCGGAGTGGACACGCTAGACCAAATGTGTTCTGTGATGACCTGCCGTAGG

The 1.4kb fragment and 750 bp TagYFP fragment (both described above) were mixed together and amplified with primer sets PstI F1 (sequence given above) and TagYFP 3R1-2:

TagYFP 3R1-2: GCGCCTGTAGCCACACCCACGGCATGGACGAGCTGTACCGGTAAAGCGGCCGCAAGAA TTCTT*GCGGCCGC*TTTACCGGTACAGCTCGTCCATGCCGTGGGTGTGG<u>CTACAG</u>GCGC

The PstI site of the TagYFP coding sequence was eliminated by replacing CTGCAG with CTGTAG to keep the same codon (cysteine) at this position. (The Not I site is in italics and the complementary sequence of the destroyed PstI site is underlined). The resulting 2.15 kb product was purified, digested with PstI and NotI and then cloned into the PstI and NotI sites of pBac[3XP3-ECFP, su(Hw)BS] ZFN-T.

The attP sequence was introduced into the plasmid as described in section 2.2.1. Subsequently, this DNA fragment was inserted into the PstI site of plasmid pBac[3XP3-TagYFP, su(Hw)BS] ZFN-T.

The blasticidin resistance gene was introduced as follows. The blasticidin resistance gene coding sequence was amplified by PCR with the primers BlasS SacII F1 and BlasS BamHI R3 from pcDNA6/V5-His-A (Invitrogen):

BlasS SacII F1: AATTACCGCGGATAAAATGGCCAAGCCTTTGTCTCAAGAAGAATCCACCCTCATT

BlasS BamHI R3: AATTGGATCCGCCCTCCCACACATAACCAGAGGGCAGCAATTCACGA

The amplified PCR product was then cloned into the SacII-BamHI site of the pIE1-3 carrying piggyBac transposase (kindly provided by Dr. Craig J. Coates, Texas A&M University). The Hr5-ie1-blasiticidin coding sequence with polyA of this clone was amplified with primer set pIE AscI F1 and pIE AscI-R1:

pIE AscI F1: ATATATGGCGCGCCCACAATCAAGTATGAGTCATAAGCTGATGTCATGTTTTGC

pIE AscI-R1: TTAATTGGCGCGCCAAGCTTAAAAGTAGGAGGAACGGGCATACTCTTGGCCACCGGCGG C

This PCR amplified product was then cloned into the AscI site of pBac[3XP3-TagYFP, su(Hw)BS] attP, ZFN-T.

### 2.3. PCR

Larvae were homogenized in Lysis Buffer (100 mM Tris-HCl (pH 8.0), 500 mM NaCl, 1% SDS, 0.15 mM spermine, 0.5 mM spermidine), and genomic DNA was isolated by phenol/chloroform and resuspended in TE buffer. All PCR was carried out using the following conditions: Step 1: 103^0^C 1 min, Step 2: 103^0^C 10 sec, Step 3: 55 ^0^C 10 sec, Step 4: 72^0^C 45 sec, Step 5: repeat steps 2-4 40 times, Step 6: 72^0^C 2 min, Step 7: 4^0^C.

#### 2.3.1. Genomic PCR for pBac[3XP3-ECFPafm] attP-ZFN-T

The 700 bp fragment from larvae transformed with pBac[3XP3-ECFPafm] attP-ZFN-T was amplified using the primers 16-1-2 F2 and 5-1-1-F2-R1:

16-1-2 F2: CAACCACTACCTGAGCACCC

5-1-1-F2-R1: GTGCAGCCAACGTCAAGCGG

#### 2.3.2. Genomic PCR for pBac[3XP3-TagYFP, hr5-ie1-BlasR, su(Hw)BS] attP, ZFN-T

Genomic PCRs of DNA from larvae transformed with pBac[3XP3-TagYFP, hr5-ie1-BlasR, su(Hw)BS] attP, ZFN-T were carried out using primer pairs TagYFP code F1 and TagYFP code R1 for PCR 1 and BlasR code F1 and BlasR code R1 for PCR 2:

TagYFP code F1: ATGGTTAGCAAAGGCGAGGAGCTGTTCGCCGGCATCGTGCCCG TagYFP code R1: CTCCAGCTTGTGGCCCAGGATGTTGCCG

BlasR code F1: ATGGCCAAGCCTTTGTCTCAAGAAGAATCCACCCTCATT BlasR code R1: TTAGCCCTCCCACACATAACCAGAGGGCA

#### 2.3.3. Inverse PCR

Inverse PCR was carried out on DNA from *Sciara* transformed with pBac[3XP3-ECFPafm] attP-ZFN-T following the Berkeley *Drosophila* Genome Project (BDGP) protocol (Roger Hoskins and Martha Evans-Holm, June 11, 2004 FlyBase http://www.fruitfly.org/about/methods/inverse.pcr.html, E. Jay Rehm, 2014 and http://www.fruitfly.org/) with minor modification: the Sau3A digested genomic DNA was cleaned with one phenol/chloroform extraction prior to the self-ligation step.

### 2.4 DNA sequencing

Amplified DNA fragments were gel purified and sequenced directly using the same primers as used for the PCR. All sequencing was done by Genewiz.

## 3. Results

### 3.1 Handling *Sciara* for transformation

To develop a transformation system, it was necessary to establish the protocols for successful injection of *Sciara* embryos (see Materials and Methods for specific details). We designed a procedure for mass mating. One day after mating, the adult female flies were impaled on agar plates and the head was crushed to induce egg laying. This is advantageous to accumulate synchronized, newly laid fertilized eggs. It is important to inject embryos before the pole cells form and while the nuclei are still in a syncytium so that the DNA injected into the cytoplasm can easily enter the nucleus. In *Sciara*, the first eleven cleavage divisions are syncytial and occur in the first nine hours at 22^0^C after egg laying (de Saint Phalle and Sullivan 1996), though the timescale seemed somewhat faster in our hands after induced egg-laying. We inject within two hours after egg-laying, which is before cleavage cycle 5 when the elimination of germ line limited L chromosomes occurs in the somatic nuclei (DuBois 1932a and 1932b and 1933; de Saint Phalle and Sullivan 1996). Also, at cleavage cycle 5, two nuclei enter the germplasm at the posterior of the egg and undergo rapid divisions to form the pole cells (DuBois 1932a and 1932b). For germline integration of DNA, it is important to inject the posterior of the embryo, and for this it is necessary to be able to distinguish the anterior and posterior ends of the embryo. The micropyle, which is a canal piercing the chorion and through which the sperm enters, is a brown disc at the anterior end of the egg (DuBois 1932a and 1932b), but is usually hard to see. Instead, we have made use of the fact that the cytoplasm is less dense at the anterior end, allowing the nuclei to be more readily visible when the embryos are on a dark background (Supplementary Figure S1). The embryos should all be aligned so that their posterior ends will be at the edge of the microscope slide for easy access for injection (Supplementary Figure S2). As a control for accuracy in determining the posterior end, the aligned embryos can be left on the agar petri dish for a week in a humid chamber to allow the black jaw at the anterior end to become visible, thus confirming the alignment. Initially, embryos were correctly aligned 90% of the time, and with practice, 99% of the time.

After embryo alignment, embryos less than 2 hours after egg-laying were used for injection. We utilized an injection apparatus that is the same for standard *Drosophila* embryo injections. However, *Sciara* embryos are extremely sensitive to environmental factors such as oxygen level, humidity, and chemical contamination. Consequently, we needed to develop a protocol to accommodate *Sciara* specific requirements using special material and handling conditions. The chorion of *Sciara* embryos is thin, and it was not necessary to remove it prior to injection. *Sciara* is extremely sensitive to the glues typically used for *Drosophila* transformation. Therefore, Kwik-Cast adhesive (World Precision Instruments, Inc) was used instead as the glue to affix the *Sciara* embryos onto microscope slides. Kwik-Cast has ultra-low toxicity and was a crucial factor for successful injection. The embryos were covered with Halocarbon oil and injected at their posterior end where the future germ cells will reside; most of the oil was removed after the injection in order to not suffocate the embryos. *Sciara coprophila* is monogenic, and females will have only sons or only daughters. We chose to inject male embryos so that the G0 male flies that emerged could be mated with five female flies, thereby amplifying the number of F1 progeny. With these procedures, we are able to inject up to 500 *Sciara* embryos/day, with a high yield of fertile adults. The non-injected embryos have 80% development to fertile adults, whereas the injected embryos had 60-70% development to fertile adults in the best cases.

### 3.2. Genomic integration of pBac[3XP3-ECFPafm] attP-ZFN-T

Prior to injections for *Sciara* piggyBac transformation, we carried out trial experiments in *Drosophila* and injected *in vitro* transcripts of mRNA encoding piggyBac transposase. However, there was much variability with each preparation of mRNA, and each had to be tested to find the maximum concentration that gave good transformation efficiency but was not too high to compromise viability and fertility of the injected embryos. Therefore, for piggyBac transformation of *Sciara*, only helper plasmid DNA encoding transposase rather than transposase mRNA was used. We utilized the *Autographa carifornica* nuclear polyhedrosis baculovirus immediate early gene (ie1) flanked by the hr5 enhancer (Mohammed and Coates 2004) instead of the hsp70 promoter (Handler and Harrell 1999) to drive piggyBac transposase; the hr5-ie1 promoter-enhancer dramatically increases transposase expression ∼500 to 1000-fold. We also constructed the same helper plasmid with a hyperactive piggyBac transposase (Yusa et al. 2011; Wellcome Trust Sanger Institute), but it did not further improve transformation efficiency in a pilot experiment done in *Drosophila*, probably because the codon usage for this transposase had been optimized for mammals rather than insects. Therefore, the standard piggyBac transposase was used for *Sciara* transformation.

The piggyBac plasmid needs to carry a dominant marker to select for positive transformants. Fluorescence markers have been widely used (Horn et al. 2002). The initial piggyBac plasmid that we used was pBac[3XP3-ECFPafm] attP-ZFN-T, which carries the ECFP selectable reporter gene (Horn and Wimmer 2000) driven by the highly conserved constitutive promoter containing three eyeless binding sites (3XP3; Noll 1993; Berghammer et al. 1999) activated by the phylogenetically conserved transcription factor Pax-6 to identify positive transformants by visualizing ECFP fluorescence of the nervous system through the larval cuticle. For future flexibility, we added the target binding site for a zinc finger nuclease (ZFN-T; Urnov et al. 2005) and also the phiC31 attP site (Groth et al. 2004), either of which could be used for site specific integration of large DNA into the ectopic piggyBac locus. We have previously reported ZFN-mediated site-specific insertion of large pieces of DNA in *Sciara* (Yamamoto et al., 2015) Based on the pilot experiments above, we injected *Sciara* embryos with (i) helper plasmid DNA encoding the hr5-ie1 driven piggyBac transposase along with (ii) a plasmid encoding the IE1 protein that binds to the hr5-ie1 enhancer to activate the promoter (Mohammad and Coates 2004), as well as (iii) the plasmid pBac[3XP3-ECFPafm] attP-ZFN-T (Figure 1A) that contains piggyBac, the selectable marker ECFP driven by the 3XP3 promoter and targeting sites of attP and ZFN for future insertion of large DNA. Out of 230 successful G0 crosses for *Sciara* transformants, we recovered two independent clones (T1 line and T2 line) which had strong ECFP expression in neural tissue, as expected from the 3XP3 promoter. Strong expression of ECFP was observed in the brain of first instar through fourth instar *Sciara* larvae (Figure 2A). From the late second instar through the fourth instar, strong expression became visible in the central ganglia on the ventral side of the larvae (Figure 2B). No non-neural ECFP signal was observed in *Sciara* larval transformants, indicating the specificity of the 3XP3 promoter. ECFP signal was not detectable in adult *Sciara* flies, probably because the black pigmentation of the adult fly body masks the ECFP signal.

**Figure 1.**
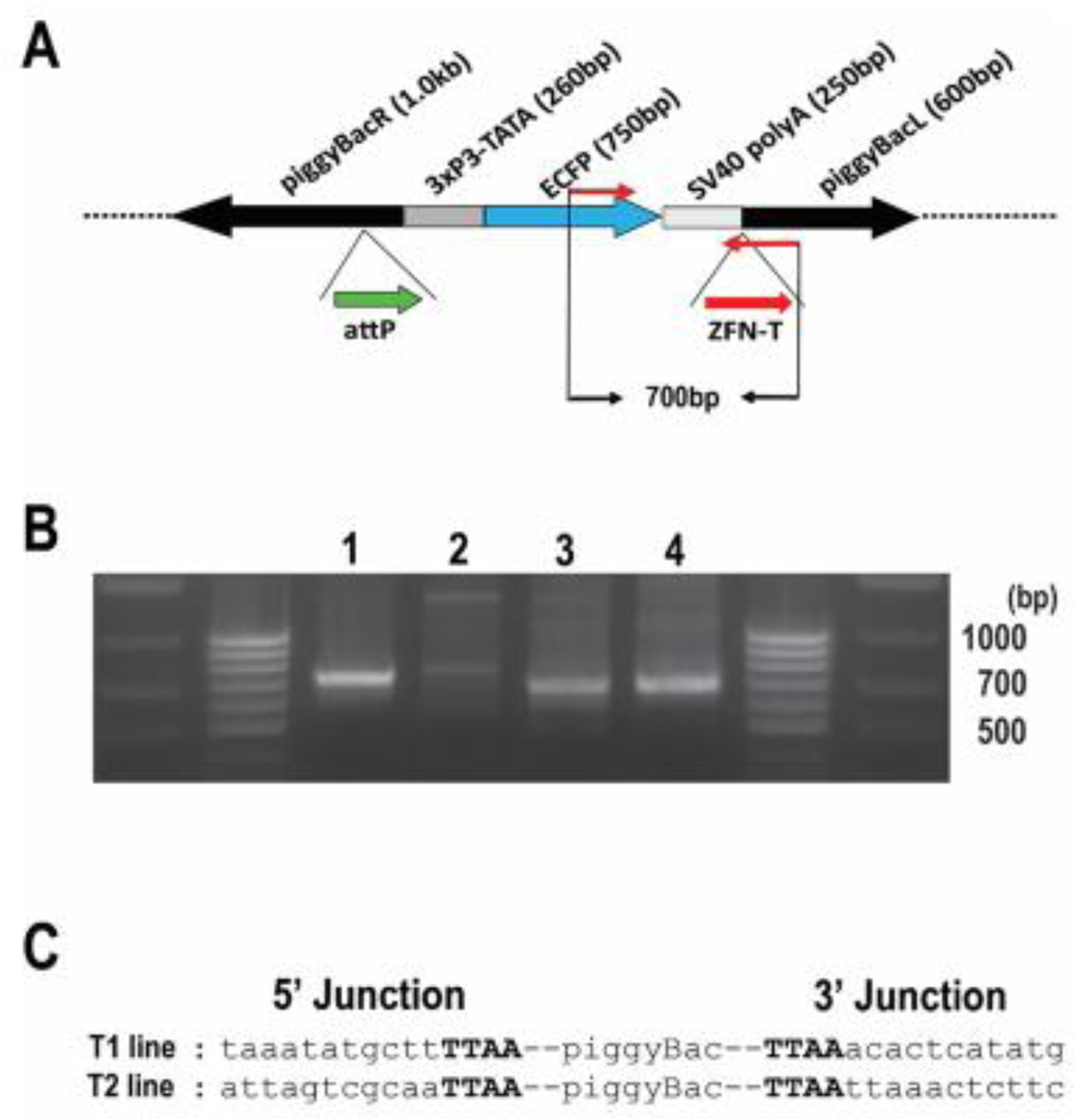
Confirmation of integration into the *Sciara* genome by piggyBac[3xP3-ECFPafm] attP-ZFN-T. **(A)** Schematic presentation of piggyBac[3xP3-ECFPafm] attP-ZFN-T. The left (L) and right (R) arms of the piggyBac vector are indicated, as is the selectable marker for ECFP driven by the 3XP3 promoter. The target sites for future insertion of large DNA are shown for phiC31 attP (attP, thick green arrow) and for a zinc finger nuclease (ZFN-T, thick red arrow). The PCR primer set to amplify a 700 bp fragment from this vector is indicated by thin red arrows. **(B)** Gel photo of genomic PCR products using internal primer pairs for the vector: (lane 1) *Drosophila* genomic DNA from larvae transformed with the same vector (positive control); (lane 2) genomic DNA from non-transformed wild type *Sciara* larvae (negative control); genomic DNA from *Sciara* transformant line T1 (lane 3) and T2 (lane 4). The 700 bp PCR product is present only in the *Drosophila* positive control (lane1), and in the two *Sciara* tranformant lines T1 and T2 (lanes 3 and 4, respectively). **(C)** Sequence of the junction between the transformation vector and genomic DNA recovered from *Sciara* genomic DNA from transformant lines T1 and T2 after using inverse PCR. The TTAA target site (in bold font) flanking the piggyBac vector is present in both *Sciara* transformant lines (T1 and T2).

**Figure 2.**
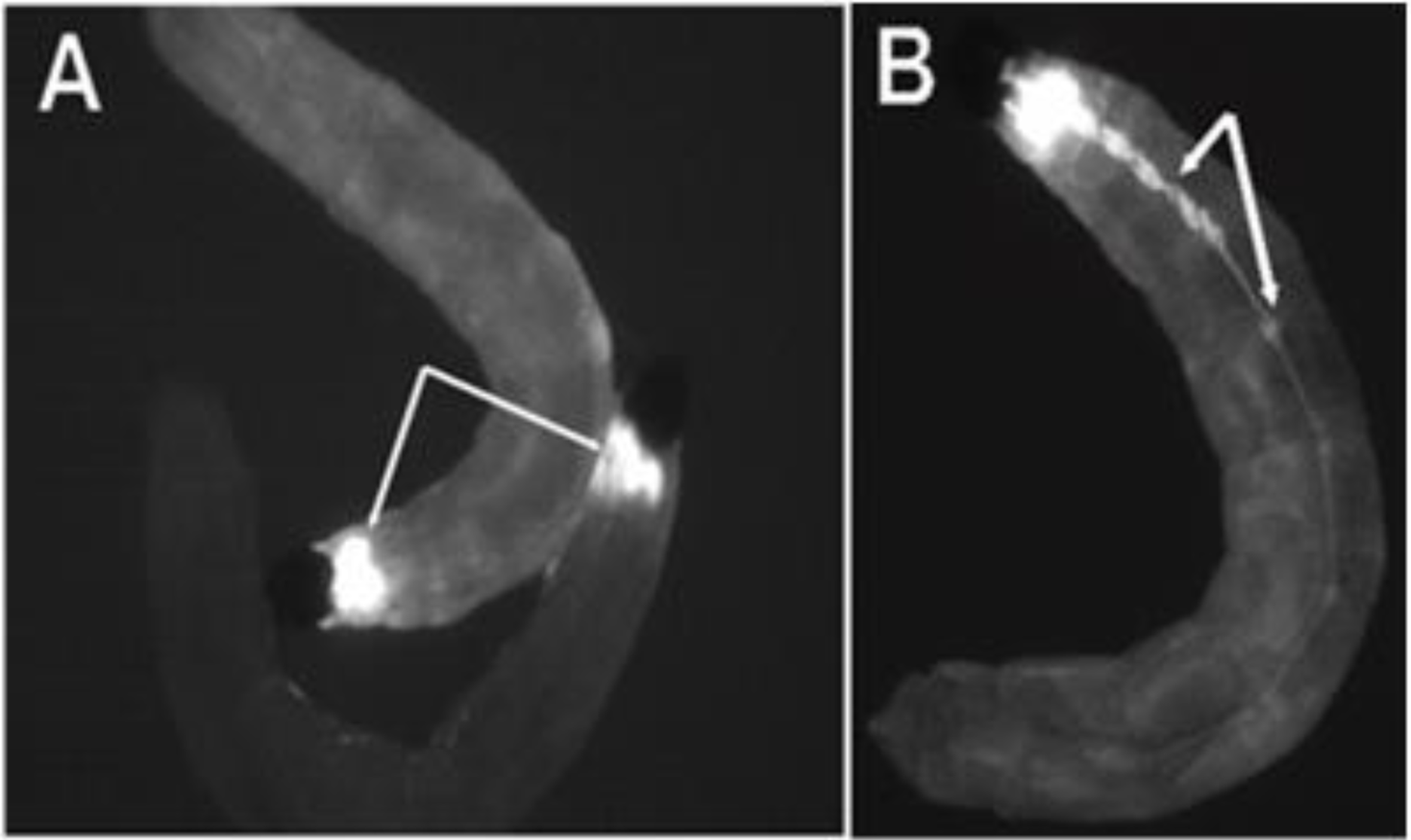
Expression of the ECFP fluorescent marker in flies transformed with pBac[3XP3-ECFPafm] attP-ZFN-T. Panels. **(A)** and **(B)** are figures of third instar *Sciara* larvae from transformant lines T1 and T2. **(A)** Strong ECFP expression is observed in the larval brain (dorsal view). **(B)** Strong expression of ECFP is observed in the central ganglion along the length of the larval body (ventral view) in addition to staining of the brain (not visualized here since that is on the dorsal side).

To confirm the integration of pBac[3XP3-ECFPafm] attP-ZFN-T vector into the *Sciara* genome, we carried out genomic PCR using a primer set to amplify a 700 bp internal sequence of the vector (Figure 1A). The same 700 bp fragment amplified from control *Drosophila* transformants was recovered from the *Sciara* transformant lines T1 and T2 but not seen in non-transformed wild type *Sciara* (Figure 1B). The flanking genomic sequences around the integrated vector was recovered by inverse PCR from *Sciara* transformant lines T1 and T2 for DNA sequencing. As is typical for piggyBac transformation, a duplicated TTAA sequence adjacent to the piggyBac inverted repeats was found in both the T1 and T2 transformant lines (Fig. 2C).

### 3.3. Genomic integration of pBac[3XP3-TagYFP, hr5-ie1-BlasR, su(Hw)BS] attP-ZFN-T

Although we have successfully recovered transformants using pBac[3XP3-EGFP afm] attP-ZFN-T, we proceeded to add some modifications to this original vector to make is more suitable for our experiments. Specifically, we **(i)** replaced ECFP with TagYFP, **(ii)** added blasticidin resistance as an additional marker for screening, and **(iii)** added the insulator element su(Hw)BS to combat position effects. The construction of this plasmid is described in the Materials and Methods and summarized in Supplementary Figure S3.

Wild-type GFP has two disadvantages for its use as a transformation marker: it is relatively insoluble and its excitation by UV light can be injurious to the transformed organisms (Horn et al. 2002). For these reasons, the more soluble blue light shifted mutant GFP variants like enhanced CFP (ECFP) have been used (Horn and Wimmer 2000; Horn et al. 2002). Nonetheless, we observed that strong expression of ECFP can have negative effects on the viability of the *Sciara* transformants. Therefore, we replaced ECFP with TagYFP (Evrogen) which is more soluble, very stable and not toxic to cells. We kept the 3XP3 promoter to drive expression of the TagYFP gene.

Though feasible, screening using just the fluorescent marker alone was extremely time consuming and labor intensive. To overcome this problem we have used antibiotic resistance as a second marker. We compared the lethality curves for neomycin (G418) to blasticidin and found that a 10X greater dose was needed for the former than the latter. Therefore, it was less costly to use blasticidin, which is routinely used for selection of cultured cells. Moreover, the blasticidin coding sequence is only 399 bp that is less than half the size of G418, thus allowing a smaller length of vector. We introduced into the piggyBac plasmid the blasticidin resistance gene driven by the hr5-ie1 promoter, and have developed the screening protocol using antibiotic selection (see Materials and Methods). After a titration experiment (Supplementary Figure S4), we chose a dose of 20 ug/ml blasticidin in the 2.2% (wt/vol) agar as optimal since this is the minimal concentration to kill all non-transformed larvae by the 4^th^ larval instar. Antibiotic resistance made it possible for us to screen a larger number of G1 progeny routinely than would be feasible with just a fluorescence marker.

We wanted to avoid position effects that not only dampen gene expression but also can interfere with *cis*-regulatory elements that drive DNA amplification (Delidakis and Kafatos 1987 and 1989). Insulator elements are DNA sequences first identified in transcription experiments (Gerasimova and Corces 2001). Insulators placed between enhancers and promoters can block their interaction; insulators placed adjacent to a transgene can protect the transgenes from chromosomal position effects on transcription. The suppressor of Hairy-wing protein binding site [su(Hw)BS] from the gypsy transposon is a strong transcriptional insulator (Geyer and Corces 1992; Corces 1995; Mallin et al. 1998; Zorin et al. 1999). The su(Hw)BS insulator also protects chorion gene locus constructs from position effects on DNA amplification in *Drosophila*, thereby allowing detailed studies on *cis*-regulatory elements and sequence elements important for DNA amplification (Lu and Tower 1997; Lu et al. 2001; Zhang and Tower 2004).

To insulate the piggyBac transgene, we introduced su(Hw)BS insulators flanking the two selection markers as shown in Figure 3A. As also noted by others (Sarkar et. al. 2006), we found that placement of su(Hw)BS in tandem as an inverted repeat (using the FseI or AscI cloning sites) without any spacer between two copies of su(Hw)BS, resulted in instability of the construct in *E. coli* during the cloning process. The su(Hw)BS inverted repeats seem to be a hot spot of transposition of the bacterial transposon Tn1000 (our unpublished data); this can be mitigated in part by using *E.coli l*acking a F’ factor. We overcame the instability problem by first putting the markers in the construct and then sequentially adding su(Hw)BS to flank the markers.

**Figure 3.**
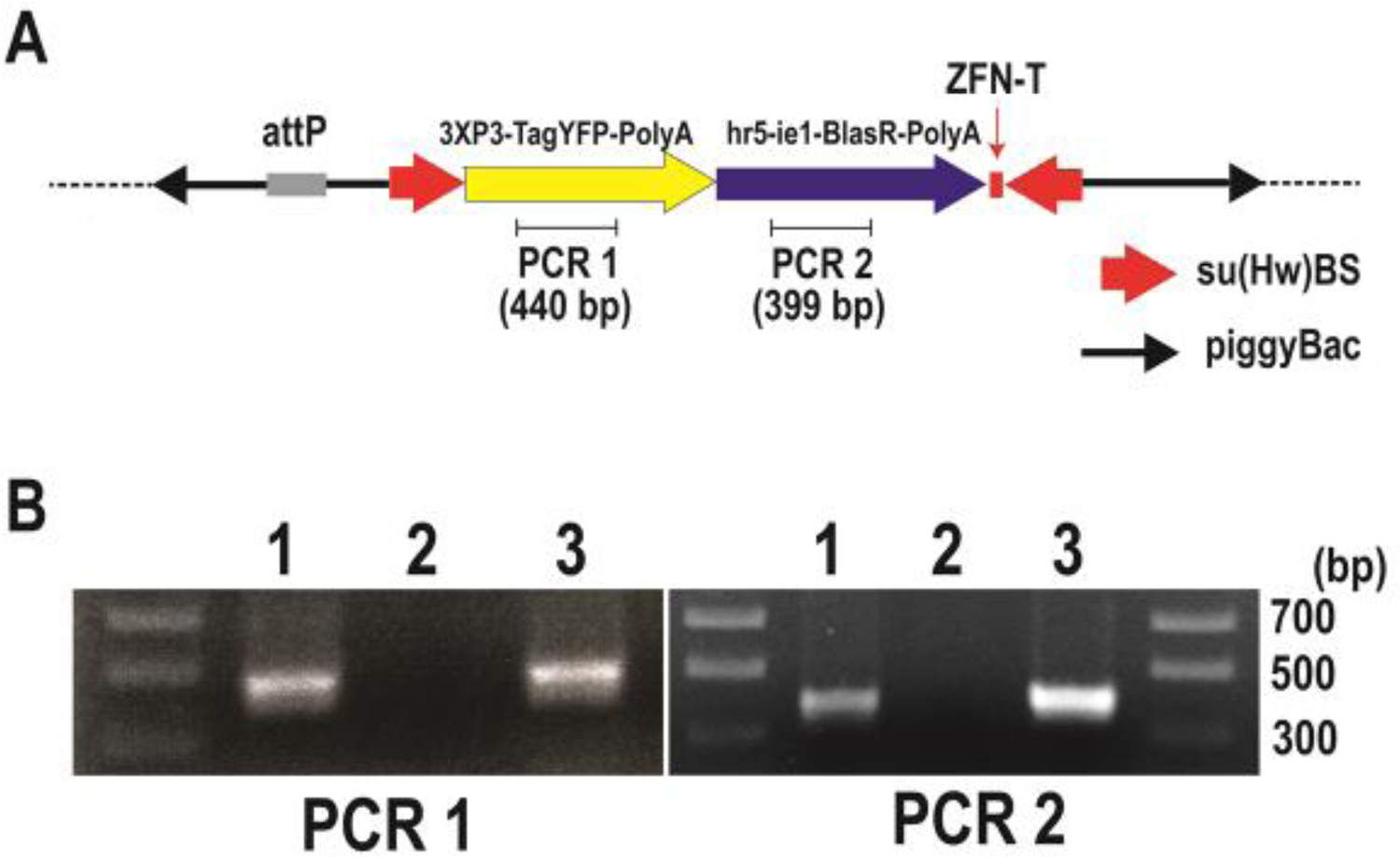
Confirmation of integration into the *Sciara* genome by pBac[3XP3-TagYFP, hr5-ie1-BlasR, su(Hw)BS] attP-ZFN-T. **(A)** Schematic depiction of pBac[3XP3-TagYFP, hr5-ie1-BlasR, su(Hw)BS] attP-ZFN-T. **(B)** Genomic PCR to amplify the TagYFP coding sequence (PCR1) and blasticidin resistance gene coding sequence (PCR2). The gel lanes contain genomic DNA from *Drosophila* transformants carrying pBac[3XP3-TagYFP, hr5-ie1-BlasR, su(Hw)BS] attP-ZFN-T (lane 1), genomic DNA from uninjected *Sciara* larva as a negative control (lane 2), and genomic DNA from *Sciara* transformant line 1 (lane 3).

We co-injected *Sciara* embryos with the same piggyBac transposase plasmid and the transactivator plasmid as used above but with the new plasmid construct pBac[3XP3-TagYFP, hr5-ie1-BlasR, su(Hw)BS] attP-ZFN-T (Figure 3A). Also, as a control, the same mixture of plasmids were injected into *Drosophila* and transformants were screened by the TagYFP marker. Among all eight independent *Drosophila* transformant lines that we recovered, TagYFP was expressed strongly in the central nervous system, eye promordium, midgut, and anal pad of third instar larvae (Figure 4, animal #1) and there was no obvious variation in the expression level of TagYFP among the lines. In contrast, variation in gut and anal pad TagYFP intensity was seen between the various *Drosophila* lines that had been transformed with the unbuffered plasmid pBac[3XP3-TagYFP, hr5-ie1-BlasR] attP-ZFN-T that lacks su(Hw)BS (Figure 4 animals #2 and #3). This suggested that the su(Hw)BS insulator was working in the eight transformant lines where it was employed. Moreover, these *Drosophila* transformants with the su(Hw)BS insulator were also resistant to the blasticidin antibiotic used as the second selectable marker. Wild type uninjected *Drosophila* larvae do not survive in the presence of 10 μg/ml of blasticidin mixed into the food, but all the *Drosophila* homozygous transformant lines survived in the presence of 80 μg/ml of blasticidin. Therefore, su(Hw)BS efficiently insulated against the position effect for hr5-ie1 driven blasticidin selection marker gene expression as well as the expression of 3XP3-TATA-TagYFP-PolyA in *Drosophila*.

**Figure 4.**
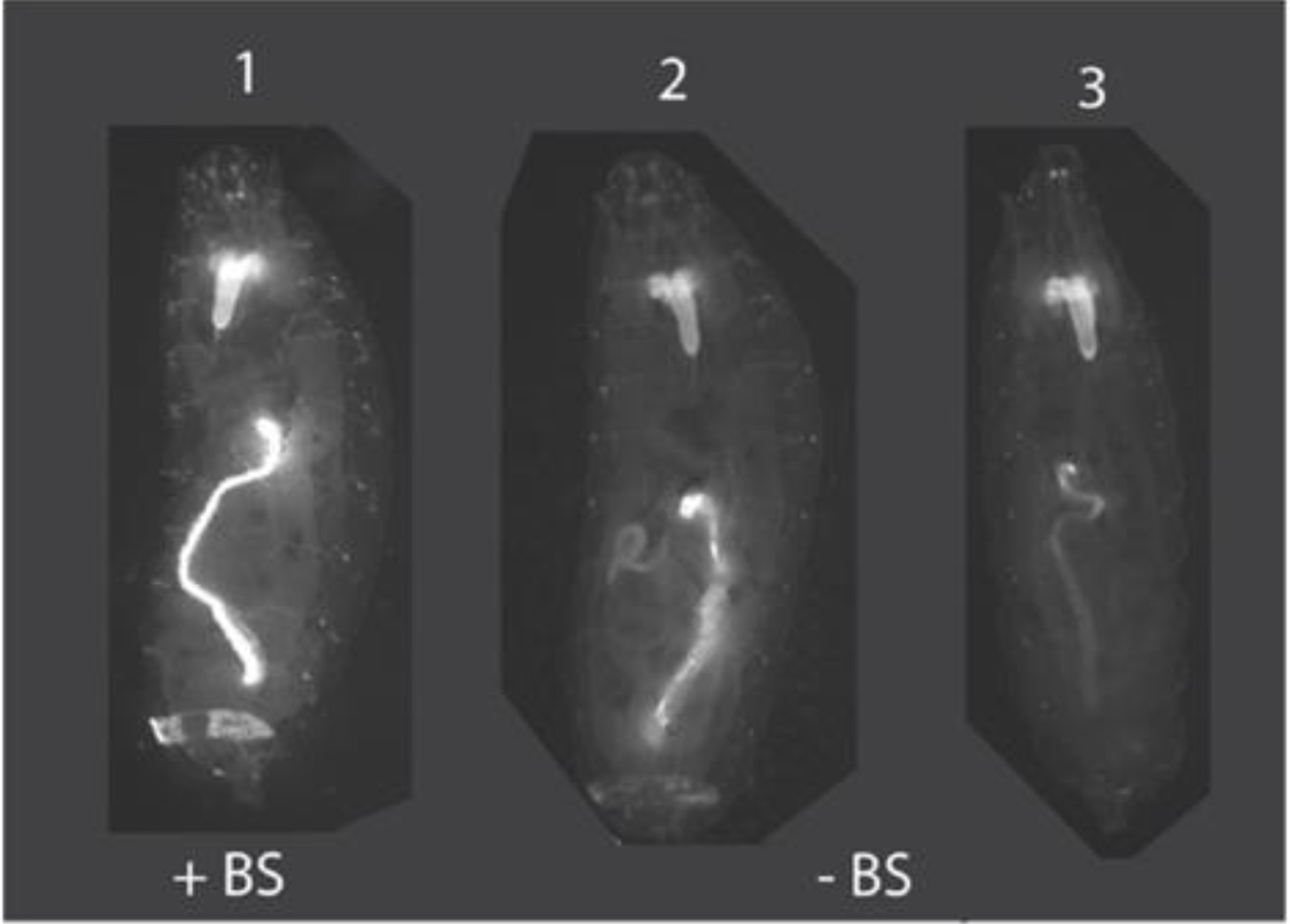
Insulation of position effects by su(Hw)BS. Animal #1 is an example of *Drosophila* transformed with pBac[3XP3-TagYFP, hr5-ie1-BlasR, su(Hw)BS] attP-ZFN-T where the su(Hw)BS insulator (“+BS”) prevents position effects and TagYFP is strongly expressed in the central nervous system, eye primordium, midgut, and anal pad of third instar larvae. This strong expression pattern with no variability was seen for all eight *Drosophila* transformant lines. In contrast, animals #2 and 3 are *Drosophila* transformed with the unbuffered construct pBac[3XP3-TagYFP, hr5-ie1-BlasR] attP-ZFN-T that lacks the su(Hw)BS insulator (“-BS”), and which show TagYFP expression in the brain in all third instar larval transformants but variable expression of TagYFP in the gut and anal pad, with expression levels differing between different transformant lines due to position effects.

After injection of *Sciara* embryos with pBac[3XP3-TagYFP, hr5-ie1-BlasR, su(Hw)BS] attP-ZFN-T, we recovered three independent blasticidin resistance lines of *Sciara* transformants (lines 1 to 3). To confirm the integration of the plasmid into the *Sciara* genome, genomic PCR was carried out using primer sets to amplify 440 bp of the TagYFP coding sequence (PCR1) and primer sets to amplify 399 bp of the blasticidin resistance gene (PCR2) (Figure 3A). The PCR products from genomic DNA of *Sciara* transformant line 1 is shown in Figure 3B. Both PCR1 and PCR2 products are present in genomic DNA from *Drosophila* transformants and *Sciara* line 1 transformants but not in genomic DNA of the negative control of uninjected *Sciara* (Figure 3B). Exactly the same results were obtained from *Sciara* transformant line 2 and line 3. The entire TagYFP marker gene was amplified by PCR and the complete TagYFP sequence was confirmed by DNA sequencing to be present in genomic DNA of all three *Sciara* transformant lines.

## 4. Discussion

We present here our systematic development of a method for transformation of the lower dipteran fly *Sciara coprophila*. The set of experiments can be used as a paradigm for development of transformation for other emerging model systems where no prior transformation methodology has existed. To begin, it was necessary to understand the biology of the organism, such as identification of the posterior end of the embryo for injection, a means to recover synchronous embryos, an understanding of when blastoderm cellularization occurs, and sensitivities to toxic conditions during handling. For example, Kwik-Cast is a very low toxic silicone elastomer developed for neuroscience applications (World Precision Instruments) The Kwik-Cast adhesive appeared to have low toxicity and enough oxygen permeability to be used for *Sciara* injection, and so far this is the only adhesive in our hands that gave a good survival rate. It could be very beneficial for other experimental organisms as well that are sensitive to environmental factors or chemicals.

We used piggyBac as the vector since it can integrate into the genomes of a wide range of insects (Handler 2002) and organisms (Lobo et al. 2006). In *Drosophila,* the piggyBac vector often integrates into the non-coding region of a gene without causing any apparent phenotype associated with it (Bellen, et al. 2011). For future flexibility, we added the target binding site for a zinc finger nuclease (ZFN-T; Urnov et al. 2005) and also the phiC31 attP site (Groth et al. 2004), either of which could be used for site specific integration of large DNA into the ectopic piggyBac locus once transformants are established.

It was also necessary to develop a method for selection of transformants. This can be challenging, especially as no prior phenotypic mutants are available for use as selectable markers. As one selection marker, we used fluorescence of the nervous system; initially, we employed ECFP and later switched to the less toxic TagYFP. In both cases, they were driven by the highly conserved constitutive promoter containing three eyeless binding sites (3XP3; Noll 1993; Berghammer et al. 1999) activated by the phylogenetically conserved transcription factor Pax-6. Although this worked, it was not feasible for scale-up for high throughput. Therefore, we also employed resistance to the antibiotic blasticidin for use in an initial screen, with subsequent confirmation by the fluorescent marker. As the promoter to drive the blasticidin resistance gene, we chose the *Autographa carifornica* nuclear polyhedrosis baculovirus immediate early gene (ie1) promoter flanked by the hr5 enhancer, since this drives 500-1000X stronger expression than other promoters (Mohammed and Coates 2004). To minimize position effects that could reduce the expression of the selectable markers, su(Hw)BS can be used as an insulator (Geyer and Corces 1992; Corces 1995; Mallin et al. 1998; Zorin et al. 1999). However, since su(Hw)BS increases the size of the vector and therefore could reduce the integration efficiency, its use is optional. Alternatively, transformants without su(Hw)BS that express the selectable markers would be indicative of insertion into the genome at sites not impacted by position effects.

**Patents:** not applicable.

## Supplementary Materials

The following supporting information can be downloaded at: www.mdpi.com/xxx/s1, Supplementary Figure S1: Visualization of the anterior end of *Sciara* embryos; Supplementary Figure S2: Schematic drawing of alignment of *Sciara* embryos; Supplementary Figure S3: construction of pBac[3XP3-TagYFP, su(Hw)BS] attP, ZFN-T; Supplementary Figure S4: Lethality curves for neomycin and blasticidin.

## Author Contributions

YY designed and executed the experiments, analyzed the results, prepared the figures and contributed to writing this paper. SAG oversaw to project and wrote the paper. All authors have read and agreed to the published version of the manuscript.

## Funding

We are grateful for research funding to SAG by NIH GM121455 currently and previously from NIH GM35929.

**Institutional Review Board:** not applicable.

**Informed Consent:** not applicable

**Data Availability:** the data are presented in this paper.

## Acknowledgments

We thank Dr. Craig J. Coates for the starting materials of the piggyBac vector, piggyBac transposase plasmid and IE1 transactivator plasmid (pIE1/153), Dr. Alfred M. Handler for piggyBac[3xP3-ECFPafm] and Dr. John Tower for the pYES vector. We also thank to Dr. Fyodor D. Urnov (Sangamo Biosciences) for kindly supplying the ZFN clones for ZFN-T. Heidi Smith and Jacob Bliss are thanked for help with *Sciara* maintenance, and Heidi Smith also for her great help in screening the transformants.

