## Supplemental Figures for "Development of transformation for genome editing of an emerging model organism"

### Supplementary Figures

**Supplementary Figure S1.** Visualization of the anterior end of *Sciara* embryos. *Sciara* embryos are 150 um X 250 µm (DuBois 1932a and 1932b). As viewed here with a dissecting microscope, the dark nuclei are more visible at the anterior end (indicated by the arrows) where the cytoplasm is less dense.

**Supplementary Figure S2.** Schematic drawing of alignment of *Sciara* embryos: **(A)** the embryos are placed on agar from a petri dish, **(B)** the edge of the agar is cut, **(C)** the embryos are aligned with the dark nuclei of the anterior end closest to the cut edge of the agar, **(D)** a microscope slide with a horizontal bar of glue (shown by green) is placed over the embryos to transfer them to the glue on the slide, **(E)** such that the posterior ends of the embryos are at the edge of the slide and accessible for injection.

**Supplementary Figure S3.** Construction of pBac[3XP3-TagYFP, su(Hw)BS] attP, ZFN-T. Steps are shown for the construction of this plasmid, as described in the Materials and Methods (section 2.2.2). Subsequently, the blasticidin resistance gene was introduced into this plasmid (not shown) to create pBac[3XP3-TagYFP, hr5-ie1-BlasR, su(Hw)BS] attP-ZFN-T.

**Supplementary Figure S4.** Lethality curves for neomycin and blasticidin. Comparison of the lethality curves for **(A)** blasticidin to **(B)** neomycin (G418). Two hundred non-transformed *Sciara* embryos were transferred to 2.2% agar petri plates with various concentrations of the antibiotic, and lethality was scored at various developmental stages.

**Supplementary Figure S1**

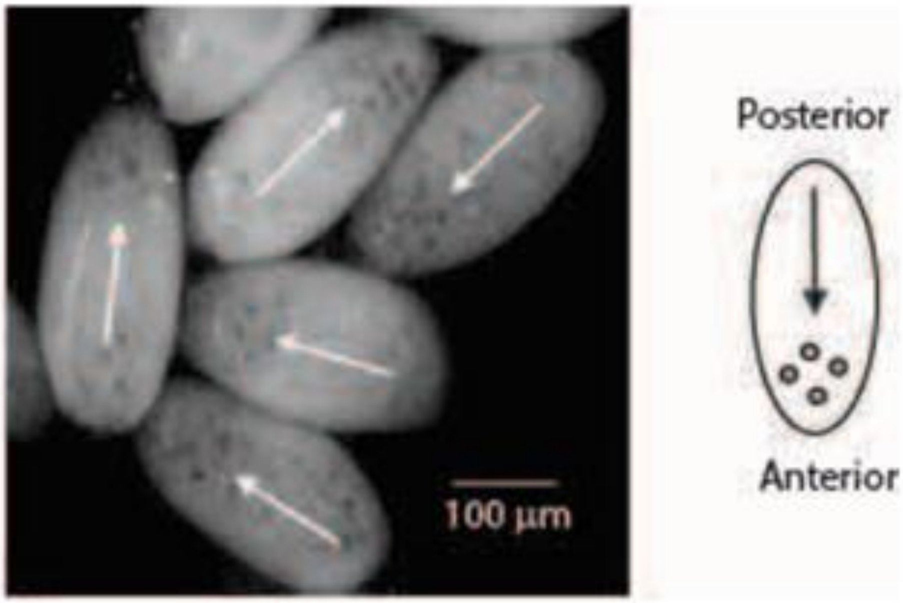

**Supplementary Figure S2**

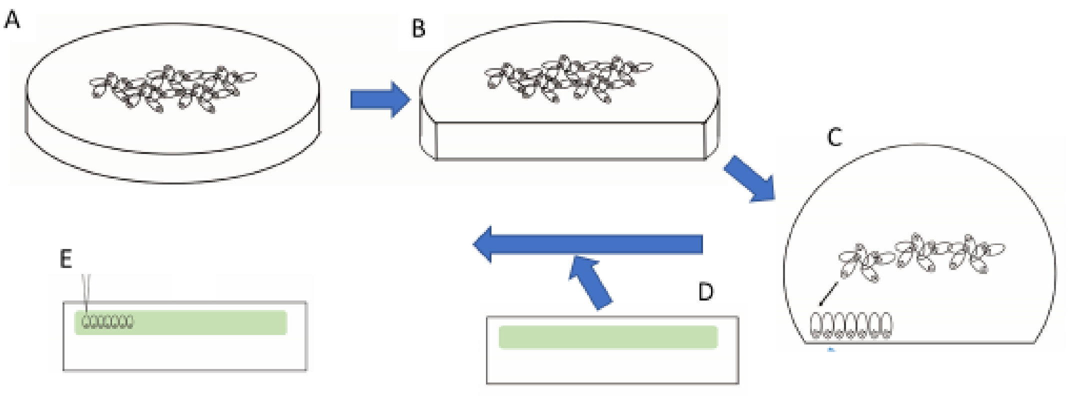

**Supplementary Figure S3**

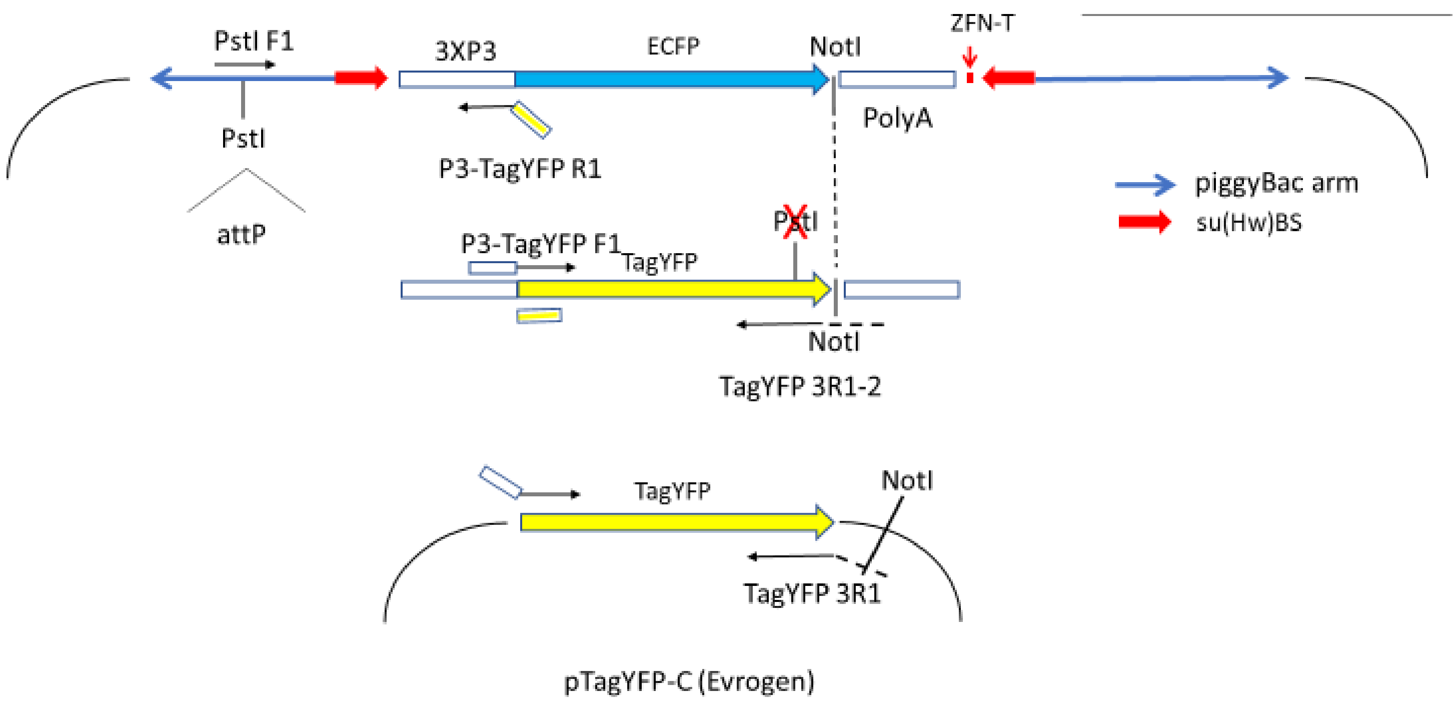

**Supplementary Figure S4**

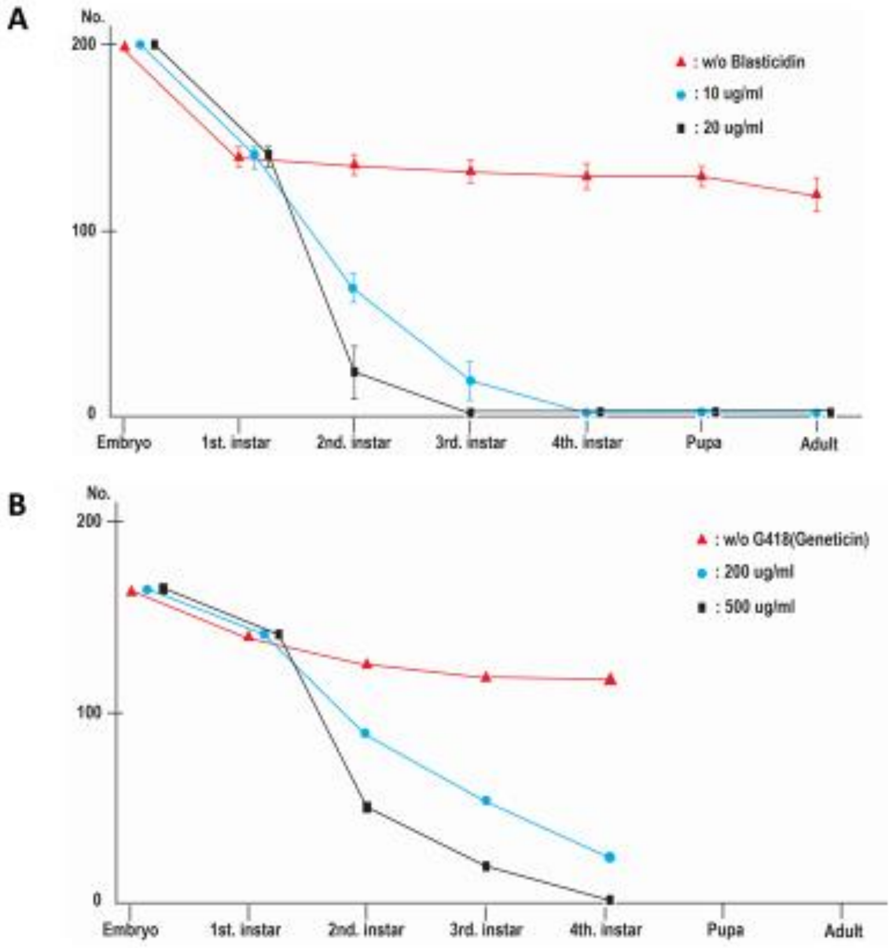
